# Neurovascular Function in a Novel Model of Experimental Atherosclerosis

**DOI:** 10.1101/2020.01.29.924936

**Authors:** Osman Shabir, Ben Pendry, Paul R Heath, Monica A Rebollar, Clare Howarth, Stephen B Wharton, Jason Berwick, Sheila E Francis

## Abstract

**Objective:** Atherosclerosis is a major risk factor for dementia. The aims of this study were to determine if experimental atherosclerosis leads to altered neurovascular function and causes neurovascular damage.

**Approach and Results:** We analysed cerebral blood volume in male C57BL6/J mice injected with an adeno-associated virus (AAV) vector for mutated proprotein convertase subtilisin/kexin type 9 (PCSK9^D377Y^) fed a Western diet for 35 weeks to induce atherosclerosis (ATH) and 9-12m male wild-type (WT) C57BL/6J. We imaged blood volume responses to sensory stimulation and vascular reactivity gas challenges in the cortex of the brain through a thinned cranial window using 2D-optical imaging spectroscopy (2D-OIS). Neural activity was also recorded with multi-channel electrodes. Stimulation-evoked cortical haemodynamics, in terms of cerebral blood volume, were significantly reduced in ATH mice compared to WT and evoked neural activity was also significantly lower. However, vascular reactivity as assessed by 10% hypercapnia, remained intact in ATH mice. Immunohistochemistry in ATH mice revealed a reduced number of cortical neurons and pericytes in the cortex, but increased astrogliosis. qRT-PCR revealed significantly enhanced TNFα & IL1β in ATH mice compared to WT as well as significant upregulation of eNOS.

**Conclusion:** Systemic atherosclerosis causes significant neurovascular decline by 9m in atherosclerotic mice characterised by reduced neural activity, associated with loss of neurons and subsequent reduced cortical haemodynamics in response to physiological stimulations. The altered neurovascular function in ATH mice is chiefly mediated by TNFα.

**Highlights:**

- Systemic atherosclerosis leads to significantly reduced stimulus-evoked hemodynamic responses in the cortex by 9m of age in the rAAV8-mPCSK9-D377Y mouse model of atherosclerosis compared to wild-type controls.
- Reduced cerebral haemodynamics are related to reduced neural activity in the cortex that could be due to a loss of cortical neurons potentially caused by significant TNFa-mediated neuroinflammation.

## Introduction

Atherosclerosis is a major risk factor for the development of dementia. The progressive atheromatous plaque build-up within cerebral arteries over time can lead to stenosis producing insufficient oxygen delivery to the brain parenchyma, potentially resulting in neuronal death and symptoms of dementia^1^. Vascular cognitive impairment (VCI) precedes the onset of dementia and may be attributed to a variety of different vascular pathologies affecting either systemic or intracranial vasculature (both large or small vessels)^2^. Due to the complexity of atherosclerosis and dementia pathogenesis, understanding the mechanisms of their mutual interactions is needed if efforts to develop therapeutics to prevent VCI and vascular dementia, which currently has no disease-modifying cure, are to succeed.

Murine models are available to study the pathogenesis of atherosclerosis and may be used to study the effects of systemic atherosclerosis on the brain. Studies using the ApoE^-/-^ knockout mouse, fed with a high fat (Western) diet, have previously demonstrated cerebral effects, including induction of neuroinflammatory changes^3^. However, a more recent virally-induced plus Western diet mouse model of experimental atherosclerosis using older mice may resemble human pathology more accurately than more aggressive models such as the ApoE^-/-^ knockout mouse model. The adeno-associated virus (AAV) vector for murine proprotein convertase subtilisin/kexin type 9 (mPCSK9) with the D377Y mutation (rAAV8-mPCSK9-D377Y) mouse model of atherosclerosis is a virally-induced gain of function mutation in the *PCSK9* gene that causes constitutively active inhibition of the LDL-receptor preventing cholesterol internalisation and degradation by hepatocytes leading to hypercholesterolaemia^4,5^. Coupled with a high-fat Western diet, the *PCSK9^D377Y^* mouse develops robust atherosclerotic lesions within 6-8 weeks with comparable LDL/cholesterol profiles to those seen in patients. In human individuals, the same gain of function mutation to *PCSK9^D374Y^* is associated with genetic hypercholesterolemia^6^.

Neurovascular coupling (NVC) ensures that active neurons; which are very metabolically demanding, are matched with an adequate and efficient blood supply (by changes to cerebral blood flow) to match the metabolic demand, facilitated by the neurovascular unit (NVU)^7^. The neurophysiological process of NVC causes localised vasodilation with a subsequent influx of oxyhaemoglobin (HbO) and decrease in deoxyhaemoglobin (HbR)^8^, which is the source of the Blood-Oxygen Level-Dependent (BOLD) signal exploited by many neuroimaging techniques. The breakdown of NVC is thought to be an important factor in the pathogenesis of many neurological conditions, including dementia^9,10^. Atherosclerosis could play an important role in the breakdown of NVC or compromise cerebrovascular function, given that it is a major risk factor for vascular dementia.

We aimed to investigate neurovascular function in middle-aged (9-12m) *PCSK9*-transduced atherosclerotic mice to address the hypothesis that systemic atherosclerosis is associated with neurovascular deficits and to identify potential mechanisms by which cerebrovascular dysfunction is induced.

## Materials and Methods

### Animals

All animal procedures were performed with approval from the UK Home Office in accordance to the guidelines and regulations of the Animal (Scientific Procedures) Act 1986, and were approved by the University of Sheffield ethical review and licensing committee. 9-12m old male wild-type (WT) C57BL/6J mice (n=5) and male C57BL/6J mice were injected i.v at 6wks with 6×10^12^ virus molecules/ml *rAAV8-mPCSK9-D377Y* (Vector Core, Chapel Hill, NC) and fed a Western diet (21% fat, 0.15% cholesterol, 0.03% cholate, 0.296% sodium; #829100, Special Diet Services UK) for 8m (ATH) (n=5). All mice were housed in a 12hr dark/light cycle at a temperature of 23C, with food and water supplied *ad-libitum*.

### Thinned Cranial Window Surgery

Mice were anaesthetised with 7ml/kg i.p. injection of fentanyl-fluanisone (Hypnorm, Vetapharm Ltd), midazolam (Hypnovel, Roche Ltd) and maintained in a surgical anaesthetic plane by inhalation of isoflurane (0.25-0.5% in 1L/min O_2_). Core body temperature was maintained at 37ଌ through use of a homeothermic blanket (Harvard Apparatus) and rectal temperature monitoring. Mice were placed in a stereotaxic frame (Kopf Instruments, US) and the bone overlying the right somatosensory cortex was thinned forming a thinned cranial optical window. A thin layer of clear cyanoacrylate glue was applied over the cranial window to reinforce the window. Dental cement was applied around the window to which a metal head-plate was chronically attached. All mice were given 3 weeks to recover before the first imaging session^11^.

### 2D-Optical Imaging Spectroscopy (2D-OIS)

2D-OIS measures changes in cortical haemodynamics: total haemoglobin (HbT), oxyhaemoglobin (HbO) and deoxyhaemoglobin (HbR) concentrations^11^. Mice were lightly sedated and placed into a stereotaxic frame. Sedation was maintained using low-levels of isoflurane (0.3-0.6%). For imaging, the right somatosensory cortex was illuminated using 4 different wavelengths of light appropriate to the absorption profiles of the differing haemoglobin states (495nm ± 31, 559nm ± 16, 575nm ± 14 & 587nm ± 9) using a Lambda DG-4 high-speed galvanometer (Sutter Instrument Company, US). A Dalsa 1M60 CCD camera was used to capture the re-emitted light from the cortical surface. All spatial images recorded from the re-emitted light underwent spectral analysis based on the path length scaling algorithm (PLSA) as described previously^11,12^, which uses a modified Beer-Lambert law with a path light correction factor converting detected attenuation from the re-emitted light with a predicted absorption value. Relative HbT, HbR and HbO concentration estimates were generated from baseline values in which the concentration of haemoglobin in the tissue was assumed to be 100μM and O_2_ saturation to be 70%.

For the stimulation experiments, whiskers were mechanically deflected for a 2s-duration and a 16s-duration at 5Hz using a plastic T-shaped stimulator which caused a 1cm deflection of the left-whisker. Each individual experiment consisted of 30 stimulation trials (for 2s) and 15 stimulation trials (for 16s) of which a mean average trial was generated after spectral analysis of 2D-OIS. Stimulations were performed in 100% O_2_ and a gas transition to medical air (21% O_2_) as well as an additional 10% CO_2_-hypercapnia test of vascular reactivity.

### Neural Electrophysiology

Simultaneous measures of neural activity alongside 2D-OIS were performed in a final acute imaging session 1-week after the 1^st^ imaging session. A small burr-hole was drilled through the skull overlying the active region (as defined by the biggest HbT changes from 2D-OIS imaging) and a 16-channel microelectrode (100μm spacing, 1.5-2.7MΩ impedance, site area 177μm^2^) (NeuroNexus Technologies, USA) was inserted into the whisker barrel cortex to a depth of ~1500μm. The microelectrode was connected to a TDT preamplifier and a TDT data acquisition device (Medusa BioAmp/RZ5, TDT, USA). Multi-unit analysis (MUA) was performed on the data. All channels were depth aligned to ensure we had twelve electrodes covering the depth of the cortex in each animal. The data was high passed filtered above 300Hz to remove all low frequency components and split into 100ms temporal bins. Within each bin any data crossing a threshold of 1.5SD above the mean baseline was counted and the results presented in the form of spikes per 100ms of data.

### Region Analysis

Analysis was performed using MATLAB (MathWorks). An automated region of interest (ROI) was selected using the stimulation data from spatial maps generated using 2D-OIS. The threshold for a pixel to be included within the ROI was set at 1.5xSD, therefore the automated ROI for each session per animal represents the area of the cortex with the largest haemodynamic response, as determined by the HbT. For each experiment, the response across all pixels within the ROI was averaged and used to generate a time-series of the haemodynamic response against time for HbT, HbO & HbR.

### Statistical Analysis

Statistical analyses were performed using GraphPad Prism v8. To compare haemodynamic & neural responses, statistical comparisons were made on HbT, HbO, HbR & MUA values using two-way repeated measures ANOVAs (with post-hoc Bonferroni correction) as well as 2-tailed unpaired t-tests. IHC and qRT-PCR data was normally distributed as per Shapiro-Wilks test and thus unpaired t-tests were used. P-values <0.05 were considered statistically significant. All the data are presented as mean values ± standard error of mean (SEM), unless otherwise stated.

### Immunohistochemistry

At the end of terminal experiments, mice were euthanized with an overdose of pentobarbital (100mg/kg, Euthatal, Merial Animal Health Ltd) and transcardially perfused with 0.9% saline and brains were dissected. One half-hemisphere of the brains were fixed in formalin and embedded in paraffin wax, with the other half snap-frozen using isopentane and stored at −80C. 5μm coronal sections were obtained using a cryostat. Immunohistochemistry was performed using an avidin-biotin complex (ABC) method (as described previously^13^). Following slide preparation and antigen retrieval, sections were incubated with 1.5% normal serum followed by incubation with the primary antibodies (rabbit monoclonal IgG anti-glial fibrillary acidic protein (GFAP) at 1:500, DAKO 1hr RT, goat monoclonal IgG anti-allograft inflammatory factor 1 (Iba1) at 1:250, Abcam 1hr RT, goat monoclonal IgG anti-platelet-derived growth factor receptor beta (PDGFRb) AF1042-SP at 1:250, R&D systems 21hrs 4C & rabbit monoclonal IgG anti-neuronal nuclei (NeuN) at 1:3000, Abcam 1hr RT). Horseradish peroxidase avidin-biotin complex (Vectastain Elite Kit, Vector Laboratories) was used to visualise antibody binding along with 3,3-diaminobenzidine tetrahydrochloride (DAB). All sections were counterstained with haematoxylin, dehydrated and mounted in DPX. Sections were imaged using a Nikon Eclipse Ni-U microscope attached to a Nikon DS-Ri1 camera. Quantifications were performed blinded using 5 sections per mouse that were chosen at random within the cortex and % coverage was quantified by computer-aided image analysis (ANALYsis^D). The mean % area was used for quantitative comparison between groups. At the end of the study period, a sibling cohort of mice were humanely killed using an overdose of anaesthetic and aortas were perfusion fixed with 10% w/v buffered formalin. *En face* Oil red O staining was performed and % lesion coverage determined by image analysis using NIS-elements software (Nikon, UK)^3^.

### qRT-PCR

Snap-frozen hemispheres were homogenised and RNA was extracted using Direct-zol RNA MiniPrep kit with TRI-reagent as per the manufacturer’s guidelines (Zymo) and RNA quality checked using NanoDrop™ (ThermoFisher Scientific). cDNA was synthesised from the extracted RNA using the UltraScript 2.0 cDNA synthesis kit (PCR BioSystems) according to the manufacturer’s guidelines. qRT-PCR was performed using PrimeTime qRT-PCR assay primers (IDT) for *IL1β*, *TNFα* and *NOS3* with *ACTB* as the reference housekeeping gene. Luna qRT-PCR Master Mix (NEB) was used with the primers, cDNA and nuclease free water and each gene for each sample was duplicated. CFX384 Real-Time System (BioRad) with a C1000 Touch Thermal Cycler (BioRad) was used to perform qRT-PCR consisting of 40 cycles. Data was analysed using the well-established delta-Ct method^14^ by normalising against *ACTB*.

## Results

### Reduced Stimulus-Evoked Cerebral Haemodynamic Responses in ATH Mice

Atherosclerosis was confirmed in ATH mice by *en face* staining of aortae with Oil Red O (0.5 vs 8.12%, n=2-4, sibling cohort). Cortical haemodynamics were recorded through the thinned cranial window using 2D-OIS whilst mechanically deflecting the whiskers of the left whisker pad. Stimulation-evoked blood volume (HbT) increase was significantly reduced in ATH-mice by 32% compared to age-matched WT controls. Oxyhaemoglobin (HbO) was also significantly reduced in ATH mice by 37% compared to WT controls, in addition to a significantly reduced washout of deoxyhaemoglobin (HbR) by 66% compared to WT controls (Figure 1A). This was irrespective of the duration of mechanical whisker stimulation (both short 2s and long 16s durations) and irrespective of the inhaled gaseous composition (100% O_2_ and 21% O_2_). Furthermore, the initial peak in 16s-stimulations; indicative of backwards dilation^15^, was also absent in ATH mice. As a final experimental condition, mice were subjected to a 10% hypercapnia test to determine vascular reactivity due to forced maximal vasodilation of all cerebral arterioles. No significant differences were observed between WT and ATH-mice (<10% difference) during hypercapnia suggesting that vascular reactivity of ATH-arterioles was not compromised (stiffened due to atherosclerosis and arteriosclerosis) (Figure 1A). The reduced cerebral haemodynamic responses in ATH mice were therefore not attributed to stiffened or narrowed cerebral vessels but potentially to a breakdown of NVC or reduced neural activity.

**Figure 1 –.**
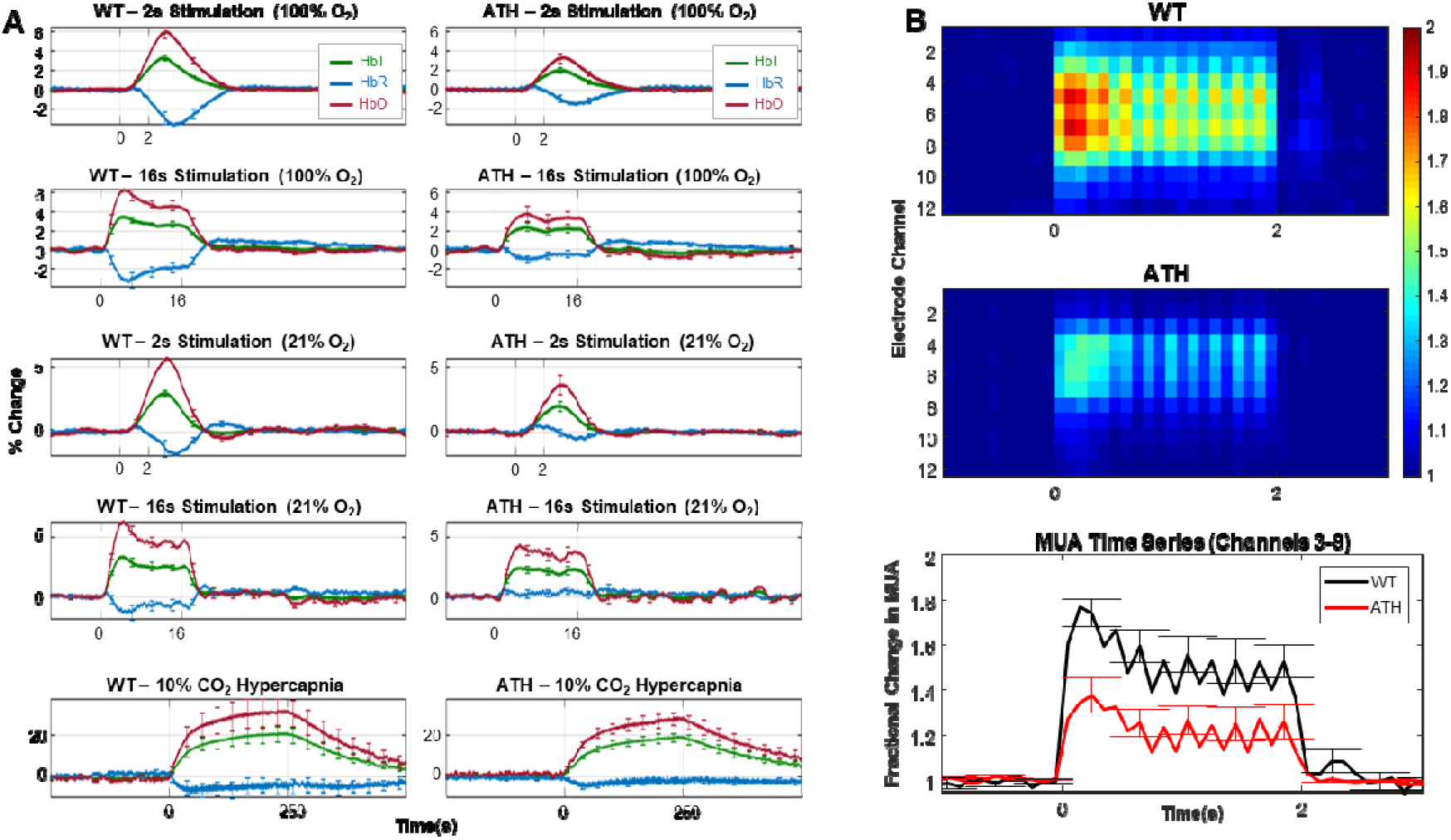
In vivo neurovascular assessments of WT and ATH mice. A) Chronic haemodynamic data to 2s/16s physiological whisker stimulations in 100% oxygen and medical air in WT (n=5) and ATH (n=5) mice. Mean HbT (area under curve (AOC) comparing WT to ATH) p=0.01 (2-way repeated measures ANOVA), HbO p=0.0005 & HbR p=0.0009. 10%-hypercapnia test of global vascular reactivity was not significantly different p=0.66 (2-tailed unpaired t-test). Peak HbT p=7.9×10^-6^, peak HbO p=4.3×10^-7^ & peak HbR p=3.98×10^-6^. B) Neural multi-unit activity (MUA) data for WT (n=4) and ATH (n=4) mice for 2s-stimulation under oxygen showing a depth map (channel depth) against duration of stimulation where somatosensory cortex is within channels 3-8. Colour bar represents fractional change in MUA. Time-series of neural MUA of WT and ATH mice with fractional changes in spikes/100ms, MUA p=0.0041 (2-way repeated measures ANOVA) for total sum AOC across all stimulations. Error bars ±SEM.

### Reduced Stimulus-Evoked Neural Activity in ATH Mice

After assessment of cerebral haemodynamics through an intact thinned skull, a final terminal experimental session was performed with an electrode inserted into the brain through a small burr-hole to record neural activity and haemodynamics simultaneously. Contrary to our expectations, neural multi-unit activity (MUA), or number of spikes/100ms, was significantly reduced by 17% in ATH-mice compared to WT controls in response to physiological stimulations across all experimental conditions and durations (Figure 1B). As such, the reduced blood volume (HbT, Figure 1) was associated with a reduced MUA and not a functional breakdown of NVC resulting in a mismatch between neural activity and blood flow.

### Neuronal Loss, Astrogliosis and Pericyte Loss in ATH Mice

After terminal experiments, mice were sacrificed, and brains were dissected for histological and genetic studies. Immunohistochemistry revealed a small but significant decrease in number of cortical neurons, as identified by nuclear NeuN immunoreactivity, in ATH mice compared to WT controls (Figure 2A). This reduction in the number of cortical neurons corresponded to the reduced neural-MUA measured in ATH mice. Staining for astrocytes revealed an increase in the number of reactive astrocytes with thicker overlapping processes within the cortex in the brains of ATH mice compared to WT (Figure 2B). Microglial numbers were not significantly altered in ATH mice, however, morphologically they displayed shorter processes compared to WT microglia (Figure 2C). Staining for pericytes revealed a decrease in pericyte coverage in ATH mice compared to WTs (Figure 2D).

**Figure 2 –.**
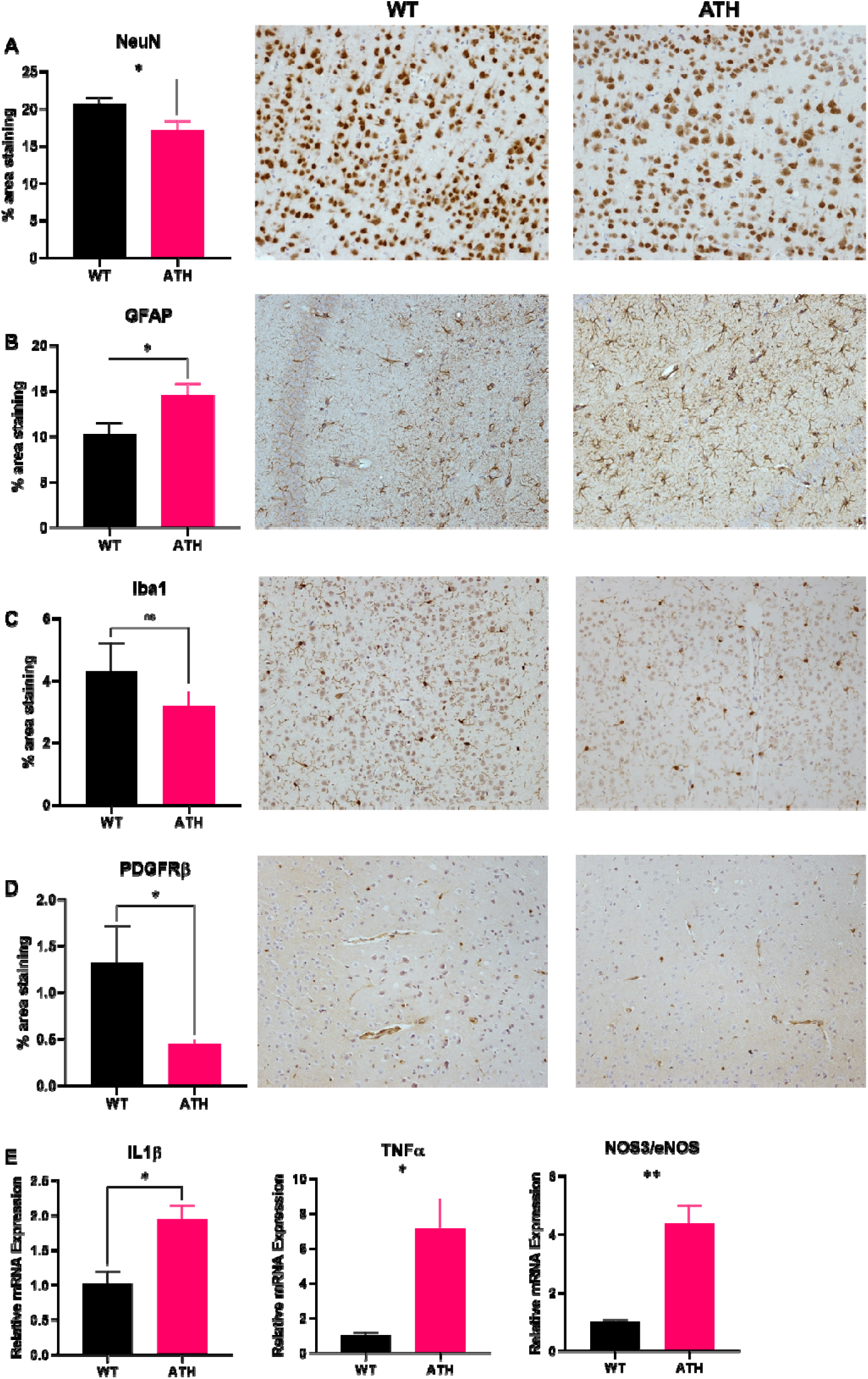
Histological and genetic assessments of WT and ATH mice. A) Mean cortical cellular quantifications between WT and ATH mice for neurons (NeuN) p=0.036 B) astrocytes (GFAP) p=0.042 C) microglia (Iba1) p=0.31 D) pericytes (PDGFRb) p=0.036. E) qRT-PCR analysis for IL1β (p=0.026), TNFα (p=0.022) & NOS3 (p=0.0045). Error bars ±SEM. Unpaired T-tests performed on all comparisons. n=4 WT and n=4 ATH.

### Significant Neuroinflammation and eNOS Upregulation in ATH Mice

qRT-PCR performed on snap-frozen brains revealed substantial upregulation of tumour necrosis factor-α (*TNFα*) by in ATH mice compared to WT controls (Figure 2E). Furthermore, interleukin-1β (*IL1β*) was also increased in ATH mice by compared to WT controls. Together these changes indicate significant neuroinflammation in the brains of middle aged ATH mice. There was also significant upregulation of *NOS3*, the gene encoding endothelial-NOS (eNOS) in ATH mice compared to WT controls.

## Discussion

The present study investigated whether systemic atherosclerosis could induce impaired neurovascular function, given that atherosclerosis is a major risk factor for dementia and the mechanisms underpinning the relationship between atherosclerosis, neurovascular decline and dementia are largely poorly understood. We found that stimulus-evoked haemodynamic responses were significantly smaller in middle-aged ATH mice compared to WT controls. The reduced blood volume in the brain could have been attributed to atherosclerosis or arteriosclerosis of cerebral vessels themselves, or potentially due to a breakdown of NVC. To address the first possibility, we tested vascular reactivity; the ability of vessels to maximally dilate in response to a 10% hypercapnia stimulus test. This revealed no significant differences between ATH and WT mice, indicating that the cerebral vessels in middle-aged ATH mice reacted to the same extent as WT vessels, and that this specific functional response is not impaired due to atherosclerosis. Intracranial arteries are more resistant to atherosclerosis and less susceptible to hypercholesterolemia^16^ than systemic arteries in humans. Intracranial atherosclerosis usually appears approximately 20 years later (usually within the 7^th^ decade of life) than systemic atherosclerosis, including coronary atherosclerosis (which tends to begin in the 5^th^ decade)^17^. The presumed reason for this is that systemic arteries tend to be elastic whereas cerebral arteries tend to be muscular with fewer elastin fibres in addition to enhanced superoxide dismutase (CuZn-SOD & Mn-SOD) levels to confer a protective antioxidant effect that is more pronounced in cerebral arteries than systemic arteries^17^. Other work in experimental atherosclerosis which studied the brain, showed Oil red O staining of extravascular lipid pools in a number of brain regions with the main phenotype being increased cerebrovascular and brain inflammation^3^.

To test whether there was a breakdown of NVC in ATH mice, neural activity was measured using a multi-channel microelectrode inserted into the active region determined from the chronic imaging sessions. Contrary to expectations, neural multi-unit activity was significantly reduced in ATH mice compared to WT controls. This implied that the cause of reduced cerebral haemodynamic responses to whisker stimulation was related to reduced neural activity within the cortex rather than a functional mismatch between neural activity and subsequent haemodynamic responses i.e. a breakdown of NVC. However, the impaired washout of HbR does suggest some level of neurovascular inefficiency and may allude to the fact neurovascular coupling is changing in ATH mice. The lack of washout of HbR under air condition suggest that although neurovascular coupling is still present, it could be less efficient resulting in inadequate oxygen delivery to active neurons. A recent study found that decreased tissue oxygenation in the LDLR^-/-^ mouse model of atherosclerosis^18^, and this is most likely to be the case in the PCSK9 model. To further assess what was causing the reduction in neural activity in the cortices of ATH mice, immunohistochemical staining for neurons was performed, which revealed a small but significant reduction in the number of cortical neurons in ATH mice compared to WT controls. This new finding indicates that systemic atherosclerosis may be able to induce neurovascular damage to the brain by causing neuronal loss. The presence of increased astrogliosis is indicative of nervous system damage, whilst upregulation of TNFα and IL1β within the brain is indicative of a neuroinflammatory response, which replicates the findings of previous research^3^.

Astrocytes are able to secrete TNFα induced by circulating cytokines, including IL1β^19^, which itself is the key inflammatory driver of systemic atherosclerosis^20^. Furthermore, a study showing the pharmacological inhibition of reactive astrogliosis by *withaferin A* was able to protect against neuronal loss by limiting the production of TNFα^21^. The *de novo* production of TNFα by glial cells within the CNS is part of the neuro-inflammatory response in the presence of neuropathology^22^. TNFα can lead to glutamate-mediated excitotoxicity as well as increased calcium-mediated excitotoxicity within neurons thus leading to a progressive neuronal loss within the brain^22^, as seen in our data. Behaviourally, it has been shown that systemic TNFα can produce cognitive dysfunction and symptoms of sickness behaviour^23^. Symptoms of delirium are also highly exacerbated by the presence of neuroinflammation, importantly due to high levels of TNFα within the brain^24^. Though we have not performed any behavioural tests on these mice, performing cognitive tests may confirm signs of early dementia and sickness behaviour in future studies. We also saw a significant increase in IL1β levels in the brains of ATH mice. It has been demonstrated that hypoxia is able to induce IL1β expression in macrophages, in addition to enhanced secretion of IL1β in the presence of cholesterol crystals^25^. However, it is important to note that our findings of neuroinflammation are related to global changes across the brain (whole homogenates) rather than being cell-specific, thus the origin of TNFα in our study cannot be determined.

We further found that eNOS levels were significantly upregulated. eNOS, releasing NO, can influence vessel tone and diameter via cGMP signalling, however, in excessive amounts, the enzyme can uncouple leading to the production of NO and O2^-^ reactive oxygen species (ROS) which subsequently react to form OONO^-^, a major cause of oxidative damage^26^. It has been demonstrated that TNFα is able to induce both eNOS in HeLa cells^27^ and iNOS in macrophages under experimental inflammatory conditions^28^. Thus, there may be a toxic vicious cycle of induction and synergy between TNFα and NOS under pathological conditions such as atherosclerosis that can contribute to neuronal death. The enhanced levels of eNOS, along with iNOS, may be a compensatory mechanism to sustain baseline vasodilation in the presence of diminished cerebral blood flow and hypoxia in the ATH mice, but also inadvertently damaging to neurons due to the potential production of ROS and interactions with TNFα. Thus, the potentially toxic combination of enhanced glial TNFα as well as eNOS may be the cause of the neurovascular deficits seen in ATH mice.

There are some notable limitations with our study. Firstly, all imaging was performed on lightly anaesthetised animals, which is known to compromise neurovascular function^29^. However, previous research from our laboratory has developed an anaesthetic regimen that is comparable to awake imaging in terms of the haemodynamic responses to physiological whisker stimulation with little effect on vascular reactivity^30^. The benefits of lightly anaesthetised preparations over awake preparations is that we can avoid the multiple considerations of behavioural state in which the animals may be whisking, grooming as well as their arousal and stress states which may be present in awake animals. Secondly, our imaging analysis assumes O_2_ saturation to be 70% with haemoglobin concentration to be 100μM, without having any direct empirical values for both. This may be important if the assumed baselines are different in the ATH animals compared to WT. Our recent study^31^ using a mouse model of Alzheimer’s disease discussed this issue in detail, which showed that regardless of the baseline blood volume estimation used, our percentage change was scaled by it (i.e. always the same change). Therefore the observation in this paper that blood volume is reduced in the ATH mice is robust.

In conclusion, we have shown for the first time that systemic atherosclerosis induces significant neural decline in the brains of middle-aged atherosclerotic mice (reduced neural MUA), associated with a mild loss of cortical neurons and astrogliosis and a reduced stimulus-evoked haemodynamic responses. Furthermore, this could also be related to synaptic damage as evidenced by neuronal loss, which precedes neuronal loss in human dementia. The neurovascular deficits seen in ATH mice may be due to the significant neuroinflammation which is evidenced by enhanced upregulation of IL1β and TNFα within the brain. Furthermore, the significant upregulation of eNOS may be a compensatory effort to maintain baseline vasodilation due to perceived hypoxia, but may also be neurotoxic due to the potential formation of OONO^-^ radicals, further contributing to and exacerbating neurovascular decline. These observations may allude to a dementia-like phenotype characterised by cognitive impairments and future work incorporating behavioural tests on these mice will allow us to establish this definitively. Future experiments probing the presence of oxidative damage, iNOS expression and apoptosis may provide a more comprehensive insight into the pathological mechanisms by which systemic atherosclerosis is able to cause such significant physiological effects on the brain and may lead to the identification of suitable therapeutic targets in the prevention of vascular mediated cognitive decline.

## Acknowledgments

We would like to thank Michael Port for building and maintaining the whisker stimulation device and 2D-OIS apparatus.

## Sources of Funding

A British Heart Foundation (BHF) project grant was awarded to Sheila E Francis to carry out the work using the PCSK9 model (PG/13/55/30365). Osman Shabir’s PhD studentship and consumables were funded by the Neuroimaging in Cardiovascular Disease (NICAD) network scholarship (University of Sheffield). Clare Howarth is funded by a Sir Henry Dale Fellowship jointly funded by the Wellcome Trust and the Royal Society (Grant Number 105586/Z/14/Z). Monica Rebollar is funded by a Conacyt scholarship.

## Disclosures

There are no competing interests to declare.

